# Genomewide basis for nitrogen use efficiency in contrasting genotypes of rice

**DOI:** 10.1101/2022.07.19.500654

**Authors:** Narendra Sharma, Dinesh Kumar Jaiswal, Supriya Kumari, Goutam Kumar Dash, Siddharth Panda, Annamalai Anandan, Nandula Raghuram

## Abstract

Rice is an ideal crop with huge germplasm diversity and post-genomic resources for improvement of nitrogen (N) use efficiency (NUE). There is a paucity of comparative studies on rice genotypes contrasting for NUE, especially with urea, the predominant fertilizer in rice growing countries. In this study, low urea-responsive transcriptomes of contrasting rice genotypes namely Nidhi (low NUE) and Panvel1 (high NUE) were compared. They were based on whole plants grown for 21 days in pots containing nutrient-depleted soil fertilized with normal (15 mM) and low urea (1.5 mM) media. There were 1497 and 2819 differentially expressed genes (DEGs) in Nidhi and Panvel1, respectively, of which 271 were common. Though 1226 DEGs were genotype-specific in Nidhi and 2548 in Panvel1, there was far higher commonality in underlying processes. High NUE is associated with the urea-responsive regulation of other nutrient transporters, miRNAs, transcription factors and better photosynthesis, water use efficiency and post translational modifications. Many of their genes co-localized to NUE QTLs on chromosomes 1, 3 and 9. Field evaluation of the contrasting genotypes under different doses of urea revealed better performance of Panvel1 in different agronomic parameters including grain yield, transport/uptake efficiencies and NUE. Comparison of our urea-based transcriptomes with our previous nitrate-based transcriptomes from the same contrasting rice genotypes revealed many common processes despite large differences in their expression profiles. Our model proposes that differential involvement of transporters and transcription factors among others contributes to better urea uptake, translocation, utilization, flower development and yield for high NUE.

**Summary:** Rice genotypes with contrasting urea use efficiency differ in the role of transporters, transcription factors, miRNAs, post-translational modifications, photosynthesis and water use efficiency

## Introduction

Sustainable nitrogen (N) management is crucial for sustainable food systems and mandated by two UN resolutions (UNEP/EA.4/Res.14 and UNEP/EA.5/Res.2), to prevent N-pollution and its impacts on health, biodiversity and climate change (Abrol et al., 2017, Sutton et al., 2019, 2020, 2021, Raghuram et al., 2021). Plant biologists have a major role in improving crop nitrogen use efficiency to minimize fertilizer wastage and pollution (Mandal et al., 2018, Raghuram and Sharma 2019, Udvardi et al., 2021, Madan et al., 2022).

Rice is among the most produced and consumed crops globally, with the lowest NUE and highest N-fertilizer consumption, making it an ideal and post-genomic crop to improve NUE. Urea is the predominant N-fertilizer used in rice-growing developing countries, whereas ammonium nitrate is preferred in the developed world. This necessitates approaching biological interventions in N-form specific manner, as was done while phenotyping for NUE (Sharma et a., 2018, 2021). Many QTLs have been identified in rice mainly for N-response and partly for NUE, though dissecting the associated genes is in progress (Waqas et al., 2018; Sinha et al., 2018, Kumari et al., 2021, He et al. 2022 and references therein).

Functional genomics revealed various N-responsive genes/processes in rice (reviewed in Kumari et al. 2021, Sharma et al., 2022, Mandal et al., 2022). Most of them were nitrate-based and only one each was based on urea (Reddy and Ulaganathan, 2015) or thiourea-based (GSE71492). Few studies have undertaken systematic shortlisting of NUE candidate genes/processes from transcriptomes associating genes with QTLs, phenotype etc. (Kumari et al., 2021). Nevertheless, some successful examples of NUE improvement in rice targeted assimilation, transporters (Hou et al., 2021), TFs (Nazish et al., 2022) and others (Chen et al., 2016, Wang et al., 2020, Khan et al., 2019, Zhao et al., 2019, Waqas et al., 2018).

Comparative transcriptomic studies focused on nitrate response in genotypes with contrasting yield (Sinha et al., 2018, Subudhi et al, 2020) or NUE in rice (Sharma et al., 2022), but such studies are lacking for urea, except the field study by Neeraja et al. (2021). In view of the difficulties with dynamic mixtures of N-forms in field soils, we report here the first controlled study of urea response in contrasting rice genotypes field validated for NUE.

## Results

### Transcriptomic analyses of rice genotypes with contrasting NUE under low urea

To understand the genome-wide effects of urea, two Indica rice genotypes Nidhi (low NUE) and Panvel1 (high NUE), were grown for 21 days (Figure 1a, 1b) in nutrient depleted soil containing normal (15 mM) and low (1.5 mM) urea and used for microarray analyses as described earlier (Sharma et al., 2018, 2021, 2022). The raw data were deposited in NCBI GEO (GSE140257). The heatmap of all samples based on Pearson correlation coefficient showed consistency in data (Figure 1c). Scatter plots revealed good correlations (R^2^) between the two independent replicates (Supplementary Figure S1) and the best two replicates were selected for further analyses. Differentially expressed genes (DEGs) were identified using geometric mean fold change value ± 1.0 (log_2_FC) with statistically significant cut off (p value ≤ 0.05). Volcano plots revealed the higher number of DEGs in Panvel1 (2819) as compared to Nidhi (1497) under low urea (Figure 1d, 1e). There were 704 up-regulated and 793 down-regulated DEGs in Nidhi, whereas 1241 up-regulated and 1578 were down-regulated DEGs in Panvel1. The DEGs common between both the genotypes were 271, of which 34 DEGs were oppositely regulated in both the genotypes (Supplementary Table S1).

**Figure 1:**
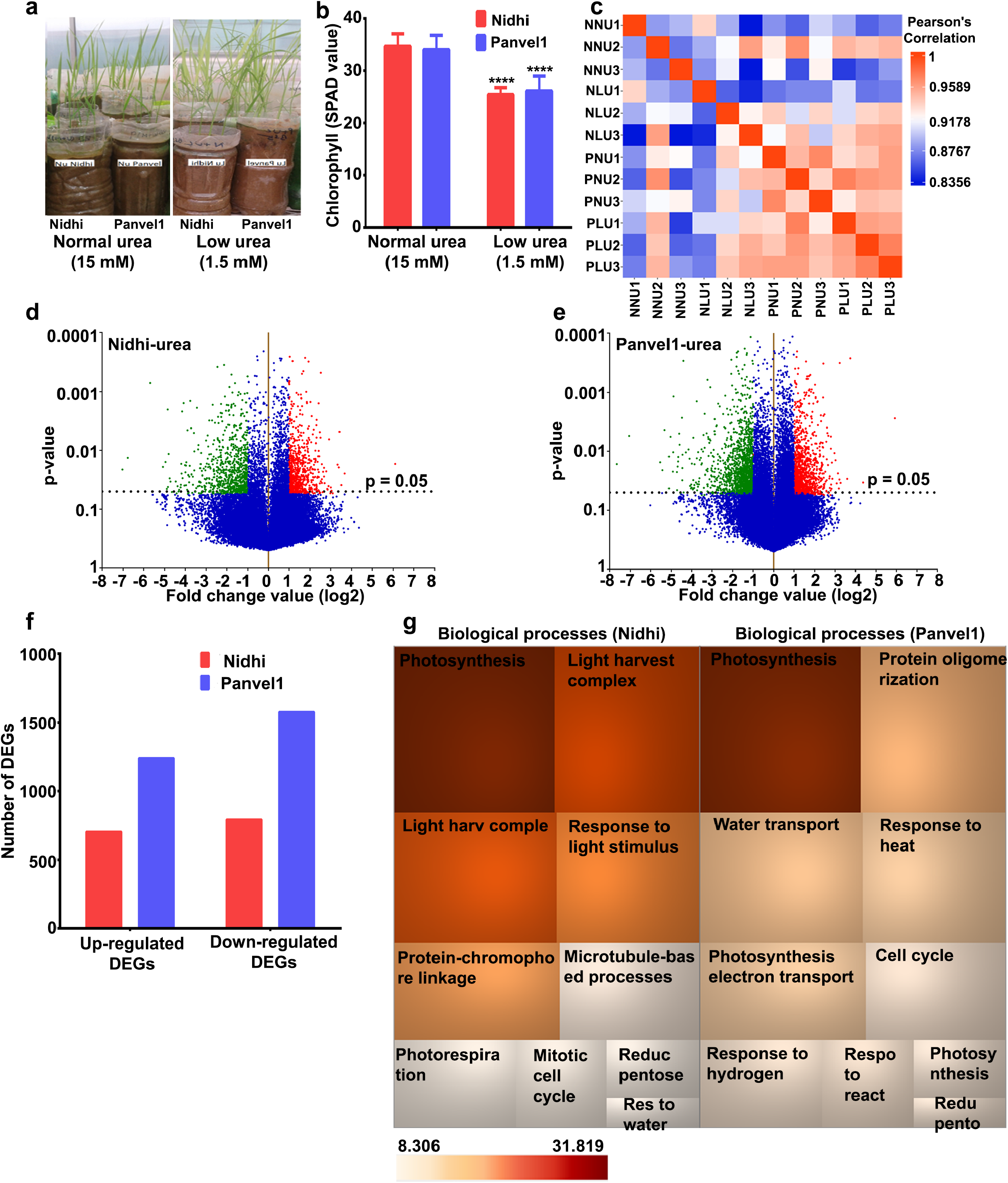
Transcriptomic analyses of low urea response in contrasting NUE genotypes. Nidhi and Panvel1 indica rice genotypes were grown in nutrient depleted soil in greenhouse under normal (15 mM) and low (1.5 mM) urea conditions (Sharma et al., 2018). Twenty-one days old grown plants (a), which showed low urea effects on chlorophyll (b) and NUE (Sharma et al., 2021) were used for microarray analyses. Unpaired T-test (b) between normal urea (control) and low urea (test) was performed in GraphPad Prism. (c) Heatmap represents the Pearson correlation coefficient (r) among the independent biological replicates of microarray data. Coloured box represents the coefficient (r) in the correlation matrix according to the scale. Urea-responsive differentially expressed transcripts are shown as volcano plots for Nidhi (d) and Panvel1 (e). Each dot on the plot represents the transcript and horizontal dashed line corresponds to p value cut-off (p = 0.05). Up-regulated transcripts are shown as red scattered dots, whereas green scattered dots represent down-regulated transcripts under low urea treatment. (f) Bar graph represents the up- or down-regulated genes detected in Nidhi and Panvel1. (g) TreeMap shows top 10 statistically significant biological processes (P < 0.05) observed in Nidhi and Panvel1. The colour of the box is according to the p value (-log_2_).

### Contrasting rice genotypes reveal common and distinct processes in response to urea

Our search for biological processes specific to the genotypes in low urea revealed a total of 353 total Gene Ontology (GO) terms for Nidhi and 493 for Panvel1. They include photosynthesis, light harvesting in photosystem I, response to light stimulus, protein-chromophore linkage, photorespiration, reductive pentose-phosphate cycle, response to water deprivation, water transport and response to cold; these biological processes are common and enriched in the urea response of both the genotypes (Figure 1f, Supplementary Table S2). Microtubule-based process, mitotic cell cycle, chromatin organization, response to oxidative stress, protein glutathionylation and alternative respiration among others were highly enriched only in Nidhi under low urea condition (Supplementary Table S2). However, biological processes related to protein oligomerization, response to heat, response to reactive oxygen species, glutathione metabolic process, cold acclimation, UDP-glucosylation and carbon fixation among others were enriched only in Panvel1 (Supplementary Table S2).

### RT-qPCR validates urea-regulation of DEGs from selected processes

Eight genes were selected based on their involvement in nitrogen transport, starch synthesis, photosynthesis, and flowering time. Their urea-regulated expressions observed in microarray data were validated by RT-qPCR under normal urea (15 mM) and low urea (1.5 mM) treatments in both the genotypes contrasting for NUE (Figure 2). Among them, four DEGs codes for Photosystem I reaction center subunit N (psaN, Os12g0189400), UDP-Glucose-dependent glycosyl transferase 703A2 (UGT703A2: Os01g0638000), days to heading on chromosome 2 (DTH2, Os06g0298200) and light-harvesting protein ASCAB9-A (ASCAB-9A, Os11g0242800) were common to both the genotypes. Except DTH2, all of them were down-regulated by low urea in both the genotypes. Among the DEGs exclusive to high NUE genotype Panvel1, three downregulated DEGs were validated viz., pseudo-response regulator 2 (PRR95, Os09g0532400), starch synthase-IIb (OsSIIb, Os02g0744700) and Nuclear Factor-Y subunit B8 (NF-YB, Os03g0413000). Among the DEGs exclusive to low NUE genotype Nidhi, an up-regulated DEG encoding a voltage-dependent anion channel 6 (OsVDAC6, Os03g0137500) was validated. The list of primers used in this study is provided in Supplementary Table S3.

**Figure 2:**
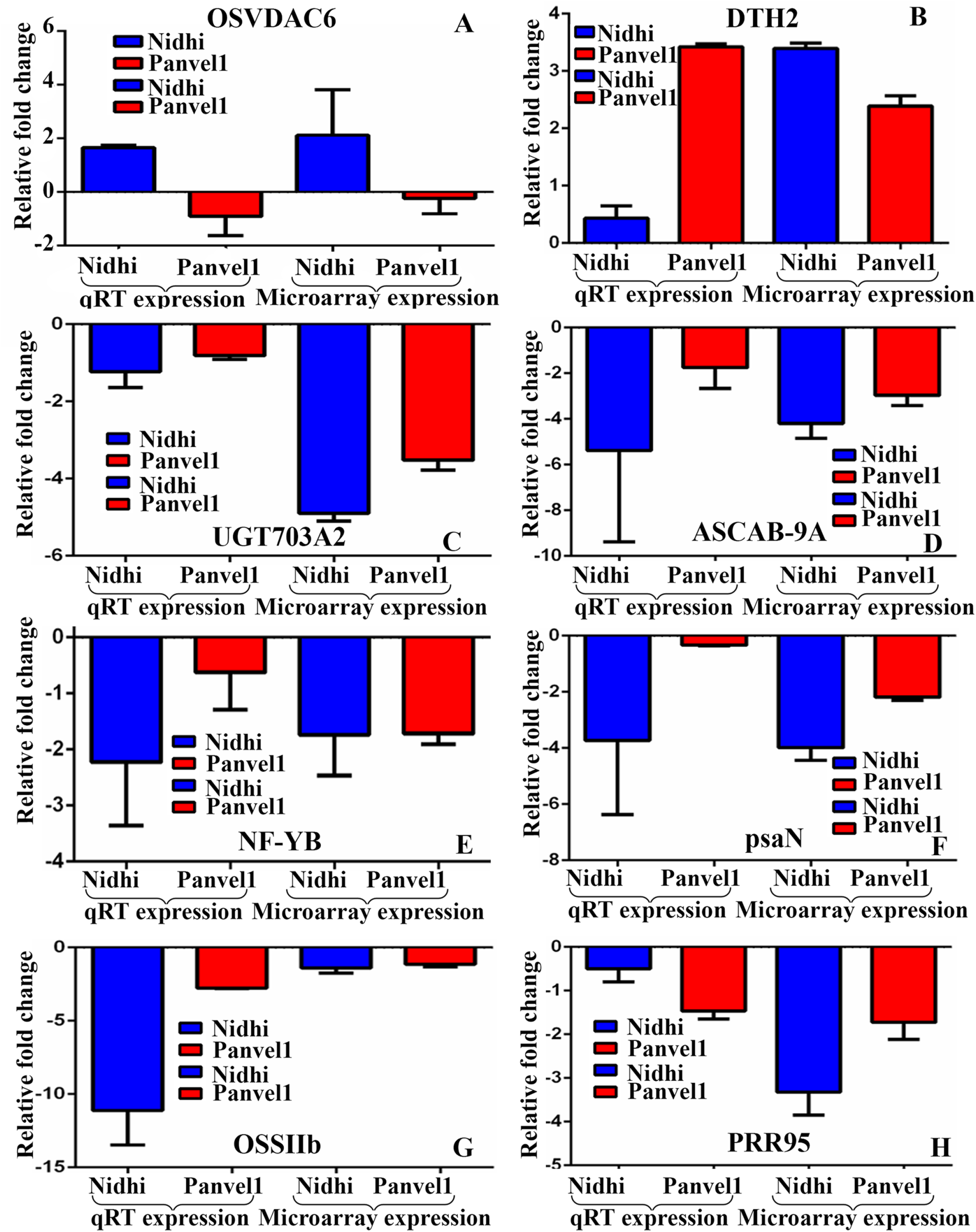
Validation of expression profile of urea responsive genes by RT-qPCR. Relative change in the gene expression was calculated by comparative Ct value method and actin gene was used for data normalization. The control values were taken as zero and the test values are shown as average of three technical and two independent biological replicates (+SE). Each sub-figure compares gene expression of qRT-PCR and microarray for Nidhi versus Panvel1 for gene OSVDAC6 (A), BZIP9 (B), UGT703A2 (C), ASCAB-9A (D), NF-YB (E), psaN (F), OSSIIb (G) and PRR95 (H).

### Transporters respond differentially to urea in contrasting rice genotypes

Nitrate transporters have been reported to regulate source-sink flux (Tegeder and Masclaux Daubresse, 2018) and NUE (Wang et al., 2018; Zhang et al. 2018; Kumari et al., 2021, Sharma et al., 2022). But urea regulation of genes encoding transporters or their roles in source-sink fluxes and/or NUE are not well understood to the best of our knowledge. In this study, mining of DEGs in transporters’ database provided 63 urea-responsive transporters belonging to 21 families for low NUE genotype Nidhi, with nearly equal proportion of up- and down-regulated DEGs (Supplementary Table S4). The high NUE genotype Panvel1 revealed 107 urea responsive transporters belonging to 36 families with more up-regulated than down-regulated transporters (Supplementary Table S4). Venn selections revealed 15 transporters common to both genotypes belonging to 10 distinct families, whereas 48 transporters were exclusive to Nidhi and 92 were exclusive to Panvel1. This clearly shows that contrasting genotypes responded differentially to urea in regulating various transporters (Supplementary Figure S2).

Further Venn selections with transporters previously linked to NUE in rice (Kumari et al., 2021) revealed 5 transporters viz. OsHKT8, OsPUP4, OsPUP7, NRAMP5 and Lsi3 as related to NUE in the genotype Nidhi. Further, 24 of the 63 transporters identified in Nidhi were associated with other physiological traits viz., abiotic stress, root and other various functions etc., whereas 35 others are completely novel and functionally unvalidated (Supplementary Table S5). Similar analysis for Panvel1 revealed 8 transporters related to NUE *viz*, OsFRDL1, MIT, OsHKT7, OsMATE2, OsNramp3, OsABCG16, NPF7.1 and OsZIP4 (Kumari et al., 2021), of which only NPF7.1 is nitrate-responsive. Twenty-six other transporters are related to abiotic stress, micro and macro nutrient regulation, water transport and development among others. Further, 73 other transporters are completely novel and functionally unvalidated (Supplementary Table S5).

Thus, we identified 13 transporters as differentially regulated by urea in contrasting rice genotypes for the first time (5 in Nidhi and 8 in Panvel1), apart from 108 novel transporters (35+73) (Supplementary Table S5). Many of them are common to urea and other N-forms to which they respond. Further characterization of these 121 urea-regulated, NUE-related transporters should enable shortlisting of candidates to improve N/urea use efficiency (UUE) in rice. Considering that barring NPF7.1, none of them is known to transport any N-form, it clearly means that NUE involves N-regulation of a large number of non-N transporters. Further, NPF7.1 seems to be an important NUE candidate, as it responds to both nitrate and urea and is also associated with yield and therefore NUE.

### Different Transcription factors mediate urea response in contrasting rice genotypes

Transcription factors (TFs) mediate plant responses to external signals by regulating the expression of the genes involved. By searching for urea-responsive DEGs in the PlantPAN3 database, we identified 45 TFs belonging to 18 TF classes in Nidhi, and 96 TFs belonging to 28 classes in Panvel1 (Supplementary Figure S2, Supplementary Table S6). In Nidhi, all the members of bZIP and Homeodomain TF families were up-regulated, while the members of AP2, C2H2, C3H and ERF TF families were down-regulated under low urea. Further, there were equal number of up and down-regulated TFs of NAC-NAM and WRKY families. In Panvel1, the members of TF family C2H2, B3, SBP, TCR, MADS and YABBY were up-regulated. Members of AP2, AT-Hook, bHLH, bZIP, ERF, homeodomain, MYB, NAC/NAM and WRKY families have more up-regulated TFs than down-regulated, while most of the HSF TF family members were down regulated than up-regulated.

To identify NUE-related TFs among these urea-regulated TFs, Venn selections was done with 237 NUE-related TFs we reported earlier (Kumari et al., 2021). It revealed, 13 NUE TFs differentially regulated by urea in Nidhi, while 18 were associated with biotic/abiotic stress, development, senescence including others, whereas 14 are completely novel and functionally unvalidated (Supplementary Table S7). Similarly, in Panvel1, 27 NUE-related TFs were found to be urea responsive (Kumari et al., 2021), while 35 TFs were linked with other traits such as abiotic/biotic stress, root, leaf, hormone, germination and phosphate regulation etc. In addition, 34 other TFs found to be urea responsive in our study are completely novel and functionally unvalidated (Supplementary Table S7). Therefore, 40 urea-regulated and NUE-related TFs (13 in Nidhi and 27 in Panvel1) should enable shortlisting of candidates for improving N/urea use efficiency (UUE) in rice, as many of them are common to urea and N-forms.

To further characterize the differential deployment of urea-responsive TFs in the contrasting genotypes Nidhi and Panvel1, TF binding sites (TFBS) were predicted/searched in the promoter regions of all DEGs using Regulatory Sequence Analysis Tools (RSAT) (http://plants.rsat.eu). A Majority of 24 DEGs in Nidhi revealed TFBS belonging to the TF family AP2EREBP, followed by 5 DEGs for NAC and 3 DEGs for bZip families, among others (Supplementary Table S8). Gene Ontology enrichment analysis using these binding sequences revealed that they are involved in translation, embryonic development ending in seed dormancy, response to cadmium ion, mismatch repair, mitochondrial transport etc., as enriched biological processes. Further, the molecular functions of genes containing these TFBS include structural constituent of ribosome, ATP-dependent helicase activity and metal ion transmembrane transporter activity in Nidhi (Supplementary Table S8). These are novel urea-responsive cis regulatory elements (CREs) unknown in any crop and therefore merit further characterization.

Similarly, in Panvel1, these TFBS are involved in protein amino acid phosphorylation, ATP binding, translation, mitochondria, embryonic development ending in seed dormancy, xanthophyll biosynthesis, mitochondrial transport etc. Their functions include protein serine/threonine kinase activity, ATP-dependent helicase activity, RNA binding CC chloroplast stroma, ATP binding etc. (Supplementary Table S8).

### Different miRNAs post-transcriptionally regulate urea response in contrasting genotypes

Post-transcriptional regulation by miRNAs is implicated in many important traits in rice development, growth, stress tolerance (Secco et al., 2013, Jiao et al., 2010), yield (Dai et al., 2018, Zhang et al., 2020) including NUE (Chao et al., 2018, Kumari et al., 2021, Sharma et al., 2022). However, they have not been characterized for urea response in NUE. Our search for miRNA targets among urea responsive DEGs using the Plant miRNA database (http://bioinformatics.cau.edu.cn/PMRD/) revealed 45 DEGs in the low NUE genotype Nidhi with nearly equal proportion of up-/down-regulated genes (Supplementary Table S9). A higher number of miRNA targets (57) were found in the high NUE genotype Panvel1, with more up-regulated than down-regulated genes as potential miRNA targets (Supplementary Table S9).

Venn selections between genotypes revealed six common miRNAs viz., osa-miRf10059-akr, osa-miR806a, osa-miRf10549-akr, osa-miR2092-5p, osa-miRf10141-akr and osa-miRf10531-akr. In order to check whether these miRNAs could significantly contribute to NUE, Venn analyses of these miRNAs was performed with 69 our recently predicted miRNA targets for NUE in rice (Kumari et al., 2021). It revealed 3 NUE miRNAs osa-miR1436, osa-miR1861c and osa-miR1863 only in Nidhi, while none found in Panvel1. Thus, our analysis in contrasting genotypes identified 42 miRNAs as novel candidates in Nidhi and 57 in Panvel1 (Supplementary Table S9) to be validated further for their role in improving NUE in rice. Genotype-wise Venn analyses of these miRNA targets with the 69 we identified earlier as NUE-related in rice (Kumari et al., 2021) yielded 3 NUE-related urea-responsive miRNAs: osa-miR1436, osa-miR1861c and osa-miR1863 only in Nidhi, while none was found in Panvel1.

Gene ontology analysis of Nidhi using ExPath 2.0 revealed their role in carbohydrate transport, metabolic process, phosphorylation and transmembrane receptor protein serine/threonine kinase signaling pathway (Supplementary Table S9). Gene ontology for target genes of Panvel1 showed processes such as cytoplasmic translational initiation, formation of translation initiation complex xylan biosynthetic process, phosphorylation, microtubule-based movement, translational initiation intracellular signal transduction and dephosphorylation. The details of their genes and functions are provided in Supplementary Table S9. These results indicate differential regulation of miRNA targets in contrasting rice genotypes and their involvement in the urea regulation of translational, post translational and carbohydrate metabolism processes, among others.

### PPI networks reveal differential urea responsive interactions in contrasting genotypes

To construct the protein-protein interaction (PPI) networks, DEG-associated interactors detected experimentally were retrieved from STING, PRIN, MCDRP and BioGRID databases as described earlier (Sharma et al., 2022). The interactors were used to construct PPI networks for each genotype and the expression values of the DEGs were mapped onto the networks in Cytoscape (Supplementary Figure S3, S4). The networks consisted of 831 nodes and 2583 edges in Nidhi, whereas 999 nodes and 3591 edges were detected in Panvel1. Venn analyses using interactors detected in both the networks revealed 317 interactors common to urea response in both genotypes, whereas 518 and 683 were exclusive to Nidhi and Panvel1, respectively (Supplementary Table 10). Their functional annotation revealed that oxidation-reduction, isoprenoid and carotene biosynthesis among others were uniquely enriched in Nidhi, whereas flower development, ovule development and regulation of meristem development were uniquely enriched in Panvel1 (Supplementary Table 10).

The complex PPI networks were sub-clustered into smaller networks (molecular complexes) using MCODE plugin in Cytoscape. In Nidhi, 11 subclusters were detected, whereas 21 subclusters were detected in Panvel1 (Supplementary Figure S5, S6). When molecular complexes with MCODE score > 4 were considered for further analyses (Supplementary Table S11), the top scorer consisted of 29 nodes and 376 edges in Nidhi, and 23 nodes and 109 edges in Panvel1. Functional annotation of genes associated with these top clusters revealed their involvement in protein translation in Nidhi and also embryo sac development in Panvel1 (Supplementary Table S11).

### Urea NUE involves differential post-translational regulation in contrasting genotypes

As post translational modifications (PTM) emerged in the GO analysis of DEGs, their details were analyzed using PTM viewer (https://www.psb.ugent.be/webtools/ptm-viewer/experiment.php). Of the 552 DEGs encoding proteins with potential PTMs in the low NUE genotype Nidhi, phosphorylation was the predominant PTM (240), followed by Lysine 2-Hydroxyisobutyrylation (224), Lysine Acetylation (52), Carbonylation (18), N-glycosylation (10), Ubiquitination (7) and Succinylation (1). The high NUE genotype Panvel1 revealed many more (852) DEGs with more extensive PTMs of their proteins in similar order, with Phosphorylation (436), Hydroxy isobutyrylation (282), Acetylation (96) and Carbonylation (27). However, it had lower Ubiquitination (6), and N-glycosylation (4) and similar Succinylation (1). Interestingly, only 118 DEGs were common to both the genotypes, clearly indicating that NUE involves targeting different proteins for PTMs in contrasting genotypes (Supplementary Table S12). Further comparison by the type of PTMs revealed that most of the differential protein targets involved lysine 2-Hydroxyisobutyrylation (59) followed by phosphorylation (49), Lysine Acetylation (9), Carbonylation (7) and Lysine Ubiquitination (1). These results highlight the hitherto unknown importance of PTMs in general and Lysine 2-Hydroxyisobutyrylation and phosphorylation in particular in urea response and NUE, which can only be identified by studying contrasting NUE genotypes.

Transporters and transcription factors are the two major functional categories in which PTMs could play crucial roles in NUE. To test this hypothesis, the 1404 urea responsive PTM targets identified here were searched among 85 transporters and 237 TFs we identified previously as NUE related (Kumari et al., 2021). It revealed 11 PTM targets as NUE related in Nidhi, including 10 transcription factors and one transporter (Supplementary Table S13). Among them, 8 TFs viz., OsPCL1, OsPRR73, RF2B, OsMyb4, OsRI, OSH15, qHD2(t) and Hd18 and a transporter (OsPUP7) were phosphorylated, while 2 other TFs, OsC3H33 and OsJMJ706 were acetylated. Similarly, the high NUE genotype Panvel1 revealed 18 TFs and two transporters as NUE related. Sixteen of these 18 TFs were phosphorylated *viz*., OsJAZ2, DOS, OsKn2, SPL6, OsIDD2, SPL16, OsSCR, YAB2, NF-YB8, OsPRR95, NF-YB10, OsTGA2, OsMYB1R1, OsMyb4, qHD2(t) and Hd18, while 2 other TFs (OsJAZ4 and OsWRKY11) were acetylated. Both the NUE-related transporters (OsHKT7 and OsNramp3) were modified by phosphorylation.

Thus, our analysis of contrasting genotypes identified 1404 urea responsive PTM targets (552 in Nidhi and 852 in Panvel1) and shortlisted 31 of them as NUE-related for the first time (11 in Nidhi and 20 in Panvel1). Further, we narrowed down 2 out of 7 types of urea-responsive PTMs as NUE-related, dominated by phosphorylation followed by acetylation. Such experimental distinction of N-response and NUE at the level of PTMs was previously unknown and could be of strategic significance for crop improvement.

### Contrasting genotypes reveal different NUE candidates by QTL co-localization

Yield-association has been our most important differentiator between N-response and NUE (Sharma et al. 2021; Kumari et al. 2021). Venn selection of 1,497 urea-responsive genes identified here in Nidhi using an updated list of 3,532 yield-related genes revealed 252 N-responsive and yield-related genes, which we termed as urea NUE-genes (Figure 3). Similar Venn selection of 2,819 urea-responsive genes in Panvel1 with 3,532 yield related genes identified 317 urea NUE genes (Figure 3). These 252 and 317 urea NUE genes were co-localized onto NUE-QTLs updated from Kumari et al. (2021). This revealed 69 unique urea NUE genes colocalized to 31 NUE-QTLs in Nidhi. They were mostly on chromosomes 1 (20), 3 (18) and 9 (9) followed by 6 each on chromosomes 5 and 7, 3 each on chromosomes 6, 11 and 12 and one on Chromosome 4 (Figure 3, Supplementary Table S14). Process annotation of these 69 urea NUE-candidates revealed carotenoid biosynthesis, starch and sucrose metabolism as the most enriched pathways in Nidhi (Supplementary Table S14).

**Figure 3:**
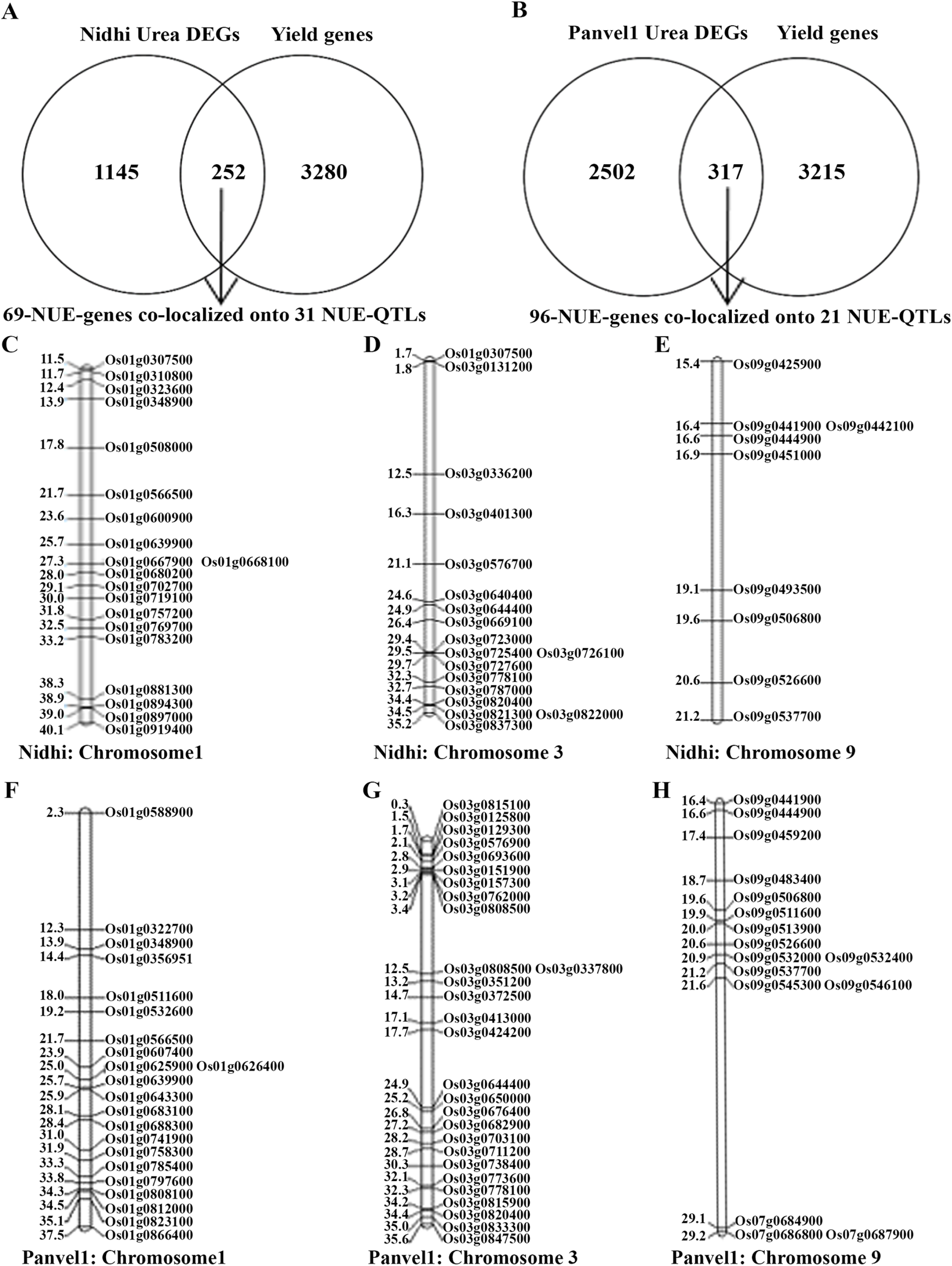
Venn selection for NUE-genes (N-responsive and yield related genes) in (1A) Nidhi urea and (1C) Panvel1 urea. Representative figures for physical location of NUE-candidates located on chromosomes 1, 3 and 9 in (1B) Nidhi urea and (1D) Panvel1 urea. Gene ID is given on the right side and the physical location of genes is given on the left side of the map (in mb).

This revealed 96 unique urea NUE genes colocalized to 21 NUE-QTLs in Panvel1. They were mostly on chromosomes 3 (28), 1 (22) and 9 (16) followed by 10 genes on chromosome 5, 6 (7), 7 (5) and 4 genes on chromosome 3 (Figure 3, Supplementary Table S14). Process annotation of these 96 urea NUE-candidates revealed circardian rhythm, cyanoamino acid metabolism, carbon fixation in photosynthetic organism, nitrogen metabolism, porphyrin and chlorophyll metabolism as the most enriched pathways in Panvel1 (Supplementary Table S14).

### Low urea enhances photosynthetic, transpiration and water use efficiencies in high NUE genotype

To experimentally test any role for carboxylation efficiency, transpiration efficiency and internal water efficiency in NUE, 21-days old potted plants of Nidhi and Panvel1 were monitored using LICOR6400XT as described in materials and methods. All three efficiencies were significantly higher (P < 0.05) in low urea (1.5 mM N) over normal urea (15 mM N) for the high NUE genotype Panvel1. Only carboxylation efficiency (but not other efficiencies) was higher in the low NUE genotype Nidhi, though to a lesser degree relative to Panvel1 (Figure 4).

**Figure 4:**
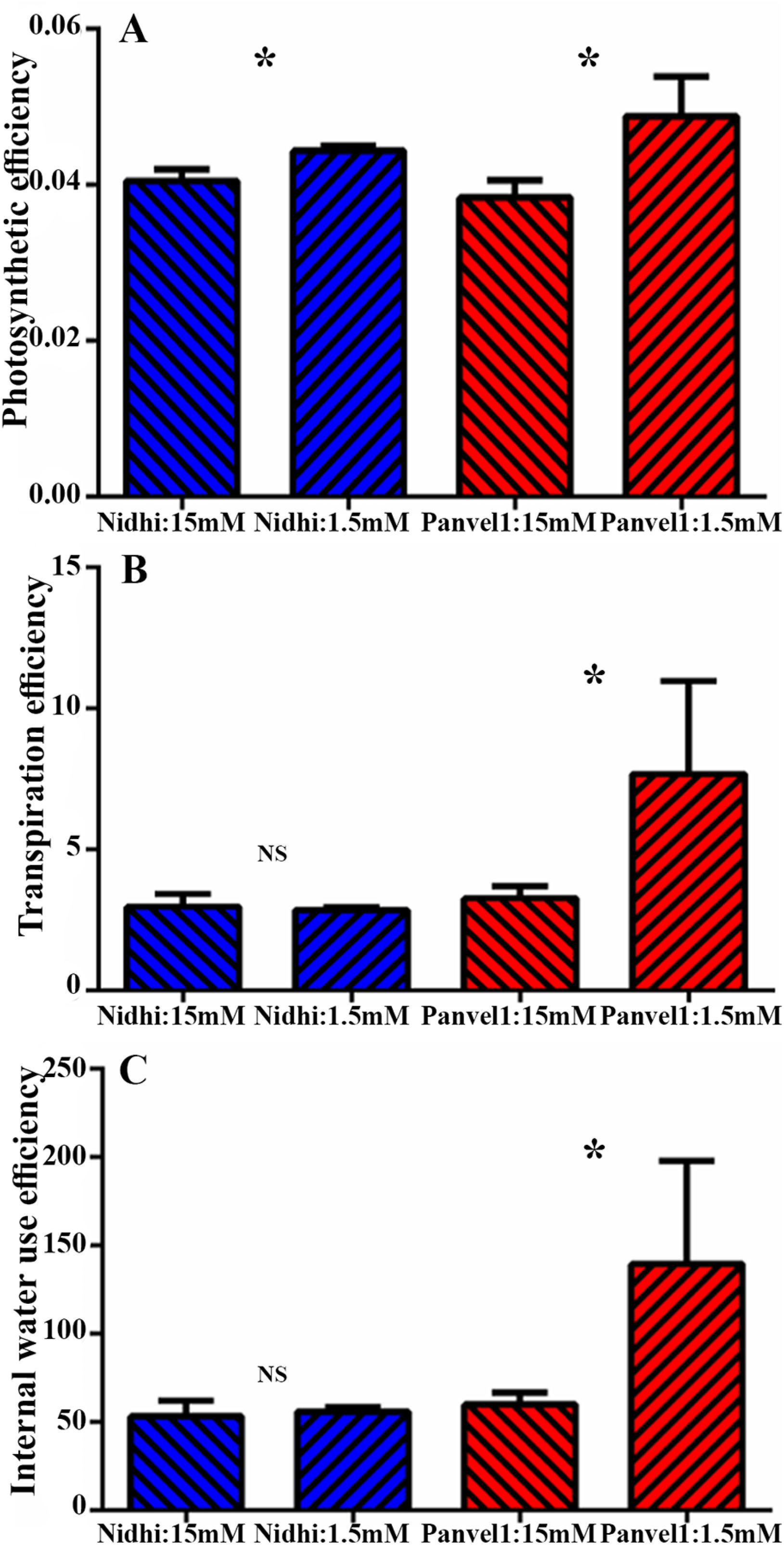
Validation of efficiencies derived associated with the biological processes: Validation was done using Licor instrument 6400XT (LI-COR, Lincoln, NE, USA) on 21 old days grown plants. Plants were grown in nutrient depleted soil and fertilized with Arnon Hoagland medium having urea as sole source of N with 15mM N concentration as control while 1.5mM, was used as test. Measurement was done in four biological replicates. Efficiencies were derived from the standard formulas as described here. (a) Photosynthetic efficiency was measured in terms of µ mol CO_2_m^-2^s^-1^/ µ mol l^-1^, (b) Transpiration efficiency was measured in terms of µ mol CO_2_m^-2^s^-1^/mmol (H_2_O)/m^-2^s^-1^ and (c) Internal water use efficiency was measured in terms of µ mol CO_2_ m^-2^s^-1^/ mol(H_2_O) m^-2^ sec^-1^). The test of significance has been shown as star (P<0.05) between the bars of low urea and normal urea, while NS represents non significance.

### Field data validate NUE in two genotypes

The field performance of the two rice genotypes with contrasting NUE used in this study (Nidhi and Panvel1) was evaluated with no added N (N0), 50 kg/ha N (N50) and 100 kg/ha N (N100) supplied as urea. The yield and NUE parameters *viz*., grain yield (kg/m2), grain yield (kg/ha), 1000 grain weight, panicle weight, grains per panicle, uptake, partial factor productivity, nitrogen transport efficiency, utilization efficiency and fertilizer N-use efficiency were measured at maturity stage. The high NUE genotype Panvel1 performed significantly better in all these parameters, relative to the low NUE genotype Nidhi (Figure 5).

**Figure 5:**
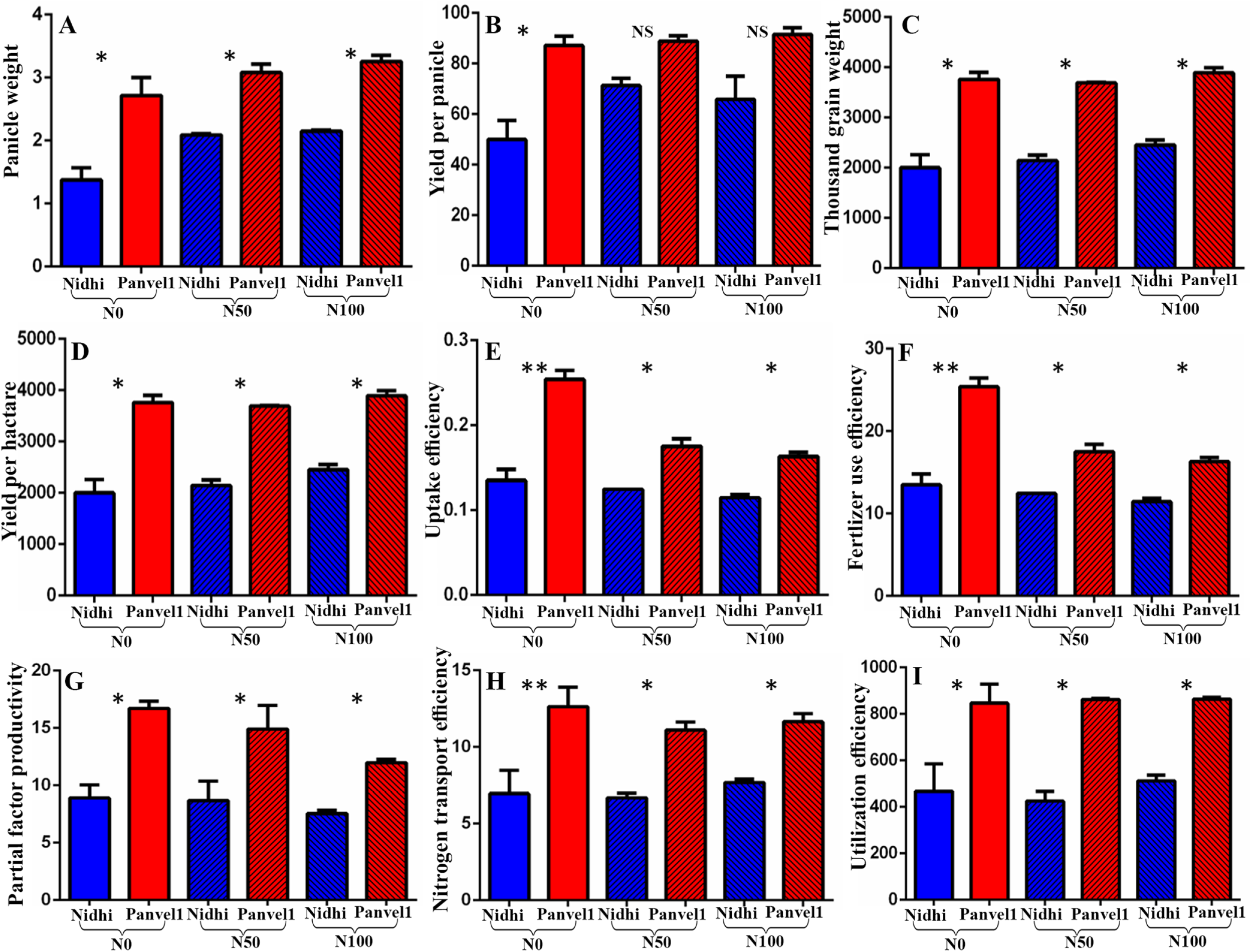
Field validation of the performance of rice genotypes: Field evaluation of the genotypes Nidhi and Panvel1 was conducted at National Rice Research Institute (NRRI-ICAR) Cuttack, Odisha, India in N0, N50 and N100 N kg of added urea/ha. Mean data of the effect of urea dose on the genotypes are shown for panicle weight, yield per panicle, 1000 grain weight, grain yield per hectare, uptake efficiency, fertilizer use efficiency, partial factor productivity, nitrogen transport efficiency and utilization efficiency. The significance levels are shown between the bars of genotype Nidhi and Panvel1 for each of the urea dose being compared (*denotes significant at P < 0.05, and ns denotes non-significant).

### Common and distinct effects of nitrate and urea on NUE in contrasting genotypes

Taking advantage of our simultaneous studies for nitrate and urea response using the same contrasting genotypes, our current urea-responsive microarray data were compared with the corresponding nitrate-responsive data (Sharma et al., 2022) (Figure 6). A higher number of 1397 DEGs responded to low nitrate in Nidhi than the 735 DEGs in Panvel1 (Figure 6a), whereas in low urea, Panvel1 displayed higher number of 2819 DEGs than the 1497 in Nidhi (Figure 6b). Venn selection of nitrate and urea for each genotype revealed fewer common DEGs in Nidhi (115) than in Panvel1 (145), suggesting that majority of the genes are unique to either nitrate or urea response in each genotype (Figure 6a). Venn analyses of DEGs assigned to the various pathways (including non-significantly enriched) revealed that a majority of these pathways are common, clearly indicating that nitrate and urea regulate N-response through the same pathways using different genes (Figure 6b, Supplementary Table S16).

**Figure 6:**
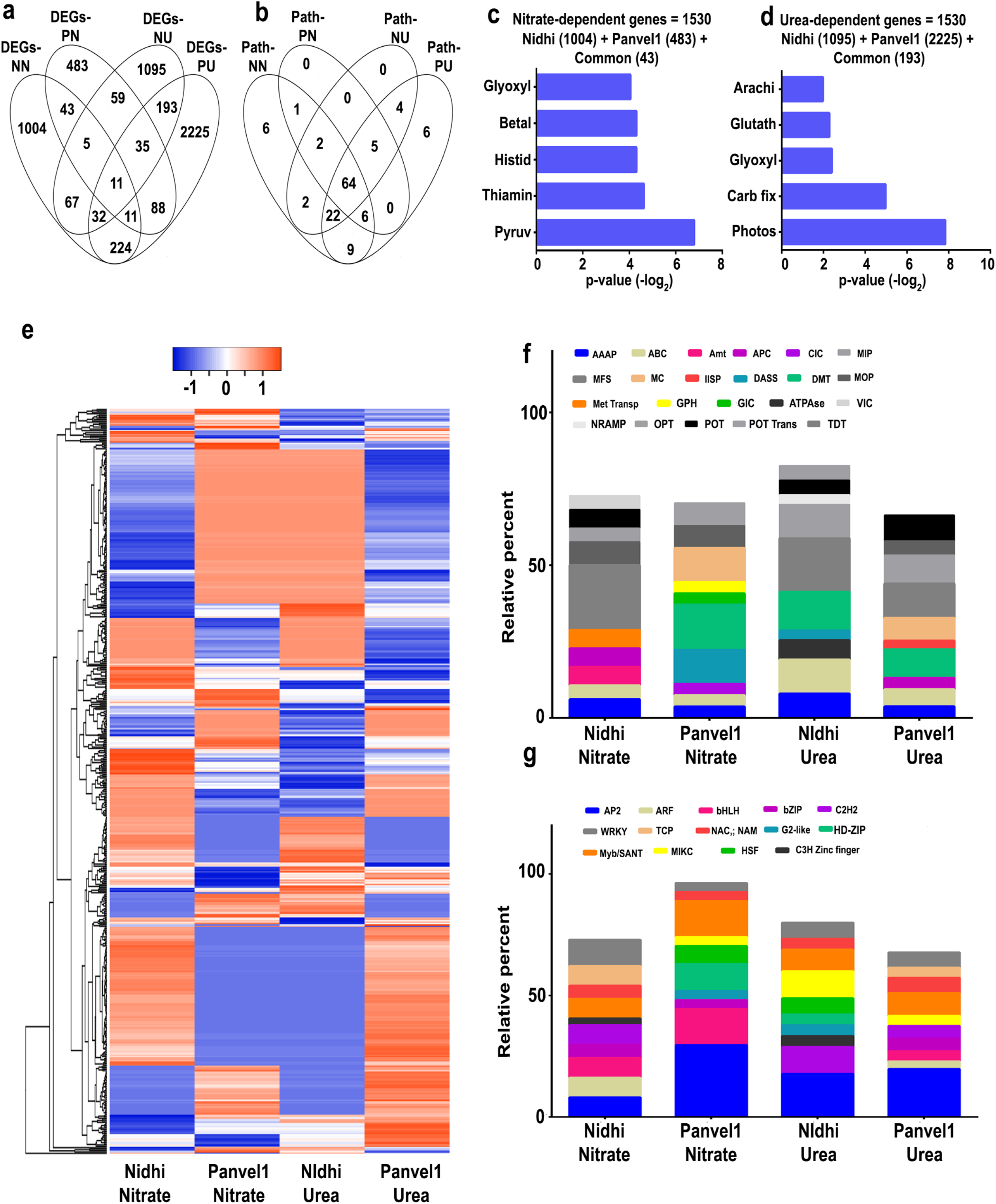
Common and specific nitrate- and urea-responsive genes and pathways in Nidhi and Panvel1. Venn diagrams represent the specific and overlap of DEGs (a) and all the assigned pathways (b) in low nitrate versus low urea responses in both the genotypes. (c) Hierarchical clustering of common N-responsive genes from Nidhi shows distinct pattern in Panvel1. Each column represents single N (nitrate or urea) treatment in a particular genotype (Nidhi or Panvel1), whereas each row represents a DEG. Stacked bar graph represents the relative percentage of top 10 enriched transporters (d) and transcription factors (e) responsive to nitrate (Sharma et al., 2022) and urea (current study). The X-axis represents the nitrate or urea treatment in Nidhi or Panvel1.

Combining the N-response data for both the genotypes revealed a total of 1530 nitrate-responsive genes and 3513 urea-responsive genes (Figure 6c and 6d). Pathway annotation of these nitrate-responsive genes revealed their involvement in pyruvate-, thiamine-, betalain- and histidine-metabolism (Figure 6c, Supplementary Table S16), Similar analysis of urea-responsive genes revealed their involved in different pathways *viz*, photosynthesis, glyoxylate and dicarboxylate- and glutathione-metabolism (Figure 6d, Supplementary Table S16). These data suggest that urea regulates carbon metabolism and associated signaling more prominently than nitrate, to modulate crop yield/NUE under low N condition. The 532 DEGs common to nitrate and urea response displayed genotype-specific patterns of expression as shown in the heatmap (Figure 6e). Evaluation of nitrate- and urea-dependent common genes revealed co-regulated, opposite and mixed expression patterns including variation in their extent of regulation for many common genes (Figure 6e).

To delineate nitrate and urea responses at the molecular level, top 10 enriched transporter families were compared, which revealed the common regulation of AAAP, ABC, MFS and POT among others under low nitrate and urea condition (Figure 6f). Enriched nitrate-regulated transporter families were Amt, ClC, GIC and GPH etc., whereas urea-regulated transporter families were ATPase, IISP and MC among others (Figure 6f). A comparison of the top 10 enriched TF families showed AP2, bHLH, bZIP, NAC and WRKY families were common to nitrate and urea response, even if individual genes are differentially regulated (Figure 6g). The TCP, MIKC and ARF among others TF families were nitrate regulated, whereas AT-Hook, ERF, and GRF were regulated by urea (Supplementary Table S16).

## Discussion

The roadmap for crop improvement towards NUE is still evolving, especially in biological terms (Udvardi et a., 2021, Raghuram et al., 2022). This is mainly due to the many different definitions of NUE (Raghuram and Sharma 2019) and inadequate experimental differentiation between N-response and NUE, especially with respect to different N-forms/doses in fertilizers and in field soils. The development of N-form and dose-specific media and nutrient-free soil aided in precise control on experimental conditions in rice (Sharma et al., 2018, 2019). They facilitated the recent characterization of the NUE phenotype and identification of contrasting genotypes validated in the field (Sharma et al., 2018, 2021), as well as transcriptomic identification of N-responsive genes/processes (Pathak et al. 2020, Mandal et al., 2022). Using yield to differentiate between N-response and NUE aided the shortlisting of important candidate genes/processes/QTLs for NUE in rice (Sharma et al., 2021, Kumari et al., 2021), including from nitrate response in contrasting genotypes (Sharma et al., 2022). The current study builds on these findings to specifically address genomewide urea response and NUE in two contrasting Indica rice genotypes for the first time. We explain their differences in NUE in terms of some common but largely differential involvement of genes/processes involving nitrogen transport, transcription factors, miRNAs, post translational modifications and QTLs. We also demonstrate differences in nitrate and urea responses and offer some candidates for crop improvement, especially since urea is the most predominant fertilizer for rice globally and NUE improvement in rice effectively means urea use efficiency.

Our transcriptomic analysis compared low urea response (1.5 mM N) relative to normal urea (15 mM N) in potted plants grown on nutrient free sterilized soil. We used the previously characterized contrasting genotypes Panvel1 (high NUE) and Nidhi (low NUE) for this purpose (Sharma et al., 2021). There were huge differences in their response to low urea, with the high NUE genotype Panvel1 showing nearly twice the number of DEGs than the low NUE genotype Nidhi, with proportionately high upregulated DEGs (Figure 1, Supplementary Table S1). This clearly indicates that high NUE may involve many more DEGs than low NUE in urea response, whereas the opposite was true in nitrate response for the same rice genotypes and conditions (Sharma et al. 2022). While these differences have been explored in detail (see later), it is important to emphasize two important findings here that were not reported in any of the many transcriptomic studies on nitrate and urea response: a) that it requires analysis of multiple (ideally contrasting) genotypes under identical conditions to discern these differences and b) all functional biological interpretations of NUE or attempts for its improvement must be made in terms of specific N-forms, or assumed to vary with other N-forms unless ruled out experimentally.

Process annotation of urea-responsive DEGs revealed photosynthesis, chloroplast and response to water (Supplementary Table S2) among those prominently regulated by urea in both contrasting rice genotypes. Photosynthesis is one of the most important processes in crop N response and NUE (Long, 2020) and its proteins are among those prominently affected by nitrogen status (Khan et al., 2022, Mu et al., 2017). Our RT-qPCR data validate the higher downregulation of the DEG PsaN (photosystem I subunit N) by low urea in the low NUE genotype relative to the high NUE genotype. These findings are consistent with similar results in wheat regarding a gene encoding photosystem II 10 kDa protein in wheat (Sultana et al., 2020). The expression of PsaN is also consistent with our physiological data showing less carboxylation efficiency in low NUE genotype Nidhi than high NUE genotype Panvel1 (Figure 6). Further, our data on higher photosynthetic efficiency, transpiration and internal water use efficiency in Panvel1 over Nidhi (Figure 6) confirm our transcriptomic findings on the involvement of these processes in urea response/NUE. They seem to be independent of N-form, as they were also reported in nitrate response/NUE in rice (Kumari et al., 2021, Sharma et al., 2022, Mandal et al., 2022).

Among the 34 urea-responsive DEGs that were common to both the genotypes but were oppositely regulated, there was Zink finger protein 15 (ZFP15). It is associated with plastid/ ion binding activity (Os03g0820400, GO:0009536) and was predicted to be involved in NUE in rice (Kumari et al., 2021). In our present study, its upregulation in the high NUE genotype Panvel1 relative to the low NUE genotype confirms its predicted involvement in NUE and makes it an attractive candidate for crop improvement.

Transporters and TFs are emerging as key regulators of N-response/NUE (Pathak et al., 2020, Nazish et al., 2021, Kumari et al., 2021, Mandal et al., 2022). Transporters exist as families for various metabolites and nutrients, including for nitrate, ammonium and urea. N-transporters are crucial for N uptake efficiency and some of them such as AMT1.1, NRT1.1A, NRT1.1B, NRT2.3b, OsNPF7. 7 (Hou et al., 2021) OsNPF7.9 (liu et al., 2022) and OsAAP1 (Yuanyuan et al., 2020) have been validated for NUE in rice. However, there are no such studies to our knowledge on urea transporters or other transporters regulated by urea in NUE in any crop. Our N-form specific study on urea response in nutrient-depleted soil devoid of urease activity clearly show that urea regulation of NUE goes beyond N-transporters found in nitrate-based studies. It is interesting that out of the 13 urea-responsive NUE related transporters we found in contrasting rice genotypes, one is a peptide transporter and others predominantly deal with other nutrients such as K, P, Zn etc. (Supplementary Table S4). Such results were also found in the SNP haplotyping of aus landraces under low nitrogen status of rice seedlings for root vigor by Anandan et al., (2022). This means urea mediates N-regulation of other transporters towards nutrient homeostasis that may be essential for nitrogen use efficiency, thus contributing to overall nutrient use efficiency. This is hitherto unknown in any crop and is worth pursuing further towards crop improvement.

Transcription Factors are not characterized in urea response but are well known in nitrate-response/NUE (Nazish et al., 2022, Mandal et al., 2018, Madan et al., 2022), though attempts to find nitrate response elements as binding sites for TFs have not borne fruit (Pathak et al., 2020). However, 7 N-responsive TFs have been validated for their role in NUE in rice (Nazish et al., 2022, Gao et al., 2020, Wang et al., 2020). In our current study, of the 128 urea-responsive TFs, 15 are known to be nitrate-responsive (Kumari et al., 2021), while the rest are novel and merit further evaluation for their role in NUE.

The role of post translational modifications (PTMs) in NUE has largely been ignored, as gene expression and protein expression dominated the literature. However, their roles in N-response, utilization, signal transduction are known (Kumari and Raghuram, 2020). Our study revealed 7 types of urea-responsive PTMs in 1286 proteins, of which phosphorylation emerged as most predominant followed by acetylation for their potential role in NUE, though ubiquitination was also found, which was linked to source-to-sink nitrate remobilization in Arabidopsis (Liu et al., 2017). More importantly, we show for the first time that high NUE is associated with more extensive PTMs in hundreds of more proteins than in the low NUE genotype. It is possible that there are many other N-regulated PTMs of proteins, whose genes are not differentially expressed in an N-responsive manner and are therefore not discernible by transcriptomics. Even proteomic comparisons of differential protein expression do not automatically reveal PTMs, unless specifically investigated for N-responsive PTMs. Future efforts to reconcile transcriptomes and proteomes for N-response/NUE must pay special attention to PTMs, as they may change the intervention strategies for crop improvement.

Many QTL for NUE have been reported in rice and the genetic dissection of associated genes is a work in progress (Kumari et al., 2021, Bai et al. 2022, He et al. 2022 and references cited therein). Our effort to co-localize urea-responsive and yield-related genes onto reported NUE-QTLs revealed that Chromosomes 1, 3 and 9 contain most of the candidate genes, adding 137 novel candidates to those reported by Kumari et al. (2021). Such chromosomal hotspots are of great value for plant breeders to improve NUE. Further, our process annotations of the genes in these chromosomal hotspots revealed their involvement in N/C fixation, porphyrin and chlorophyll metabolism etc., in high NUE and carotenoid, carbon and sucrose metabolism in low NUE genotype.

Our comparison of the present genomewide urea response data with our previous nitrate-response data generated in the same genotypes under identical conditions provides three important insights that can only be obtained in such comparisons. First, among the well-known common DEGs between nitrate and urea, a urea transporter (OsDUR3) was downregulated by low nitrate but upregulated by low urea conditions. Second, among transporters, ammonium transporters were regulated primarily by nitrate, whereas ATP-dependent ion transporters were regulated by urea, clearly indicating mutual regulation of the uptake/transport of different N-forms that commonly exist in field soils. Third, TFs families such as TCP were nitrate-responsive, while AT-hook TF was urea-responsive (Supplementary Table S6). It has been shown that AT-hook TF interacts with TCP TF (Yin et al., 2018), which was involved in the regulation of nitrate response in rice (Guan et al., 2017). While their role in rice NUE needs validation, our results indicate that their regulation by different N-forms may play a role in it as a means of balancing the plant’s needs in response to variable mixtures of different N-forms in field soils.

We propose a hypothetical model on the possible role of urea-responsive DEGs in NUE, (Figure 7), which broadly classified into two categories viz. those DEGs and associated processes/pathways enriched in high NUE genotype, while others enriched in low NUE genotype. We observed that in response to urea, distinct set of transporters and TFs are differentially enriched in contrasting genotype. For example, OsPIN1d (Auxin Efflux Carrier (AEC) Family: Os12g0133800) was not detected in Nidhi, whereas it was upregulated in Panvel1 (Supplementary Table S1). It has been shown that rice plant knocked out for OsPIN1d along with OsPIN1c gives decreased plant height, tiller number and severely disrupted panicle development (Jiajun et al., 2022). Our data shows its’ up regulation in Panvel1 could be linked to high NUE of Panvel1 over Nidhi. It has been shown that OsGRF4 transcription factor promotes hormone-dependent NUE enhancement in rice (Li et al., 2018). Most of the DEGs from GRAS family transcription factors were up-regulated in Panvel1, whereas they were not detected in Nidhi under low urea condition (Supplementary Table S4). This suggests their role for efficient NUE in Panvel1. Finally, our field evaluation of these two rice genotypes namely Nidhi and Panvel1 contrasting for NUE, clearly reveals that high yield and nitrogen use efficiency of genotype Panvel1 over the Nidhi is due to better nitrogen uptake, better N-transport and better N-utilization (Figure 7).

**Figure 7:**
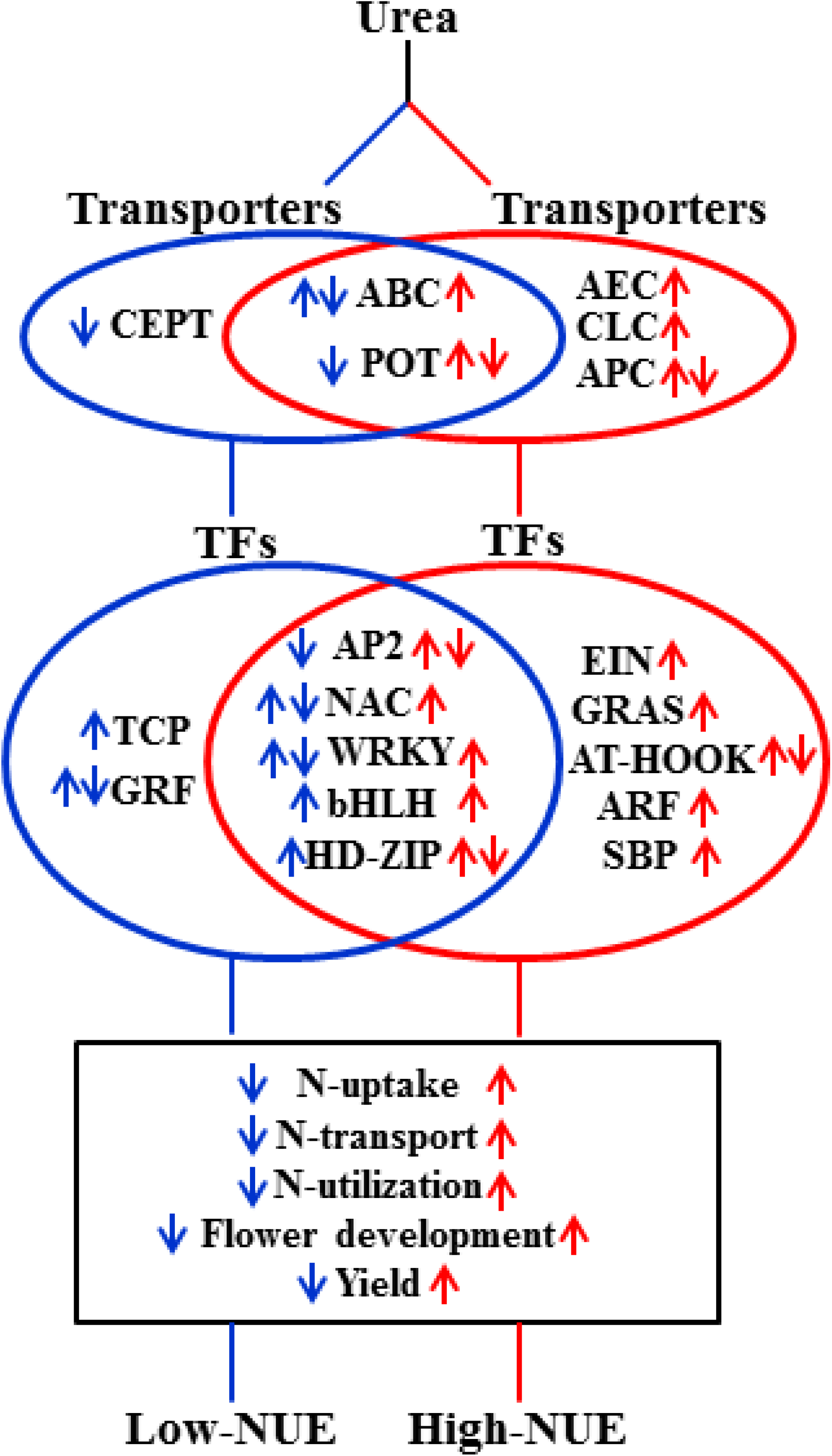
Hypothetical model depicting the important classes of genes and associated pathways differentially regulated in low-(Nidhi) and high-(Panvel1) NUE genotypes. Blue and red colour denote the differential regulation of gene classes/pathways in low- and high-NUE genotypes, respectively. Up- and down-ward arrows represent the upregulation and downregulation, respectively in a particular class/pathways. Both the arrows together represent mixed regulation of gene classes/pathways. TFs, transcription factors.

## Materials and Methods

### Plant material, growth conditions, N-treatments and physiological measurements

Two genotypes of rice (*Oryza sativa* ssp. *Indica*) namely Nidhi and Panvel1 were chosen, based on contrasting germination, yield and NUE (Sharma et al., 2018, Sharma et al., 2021). Their seeds of modal weight were surface-sterilized and grown in pots containing nutrient-depleted soil exactly as described (Sharma et al., 2018, Sharma et al., 2019). The pots were saturated with modified Arnon-Hoagland medium containing urea as the sole N source at 15 mM (normal) or 1.5 mM (low) concentration as control and test conditions as described earlier (Sharma et al. 2021). The pots were replenished with media to saturation every few days and plants were grown for 21 days in the green house at 28 **°**C and 70% relative humidity with 270 μmol m^-2^ s^−1^ light intensity and 12/12 hr photoperiod. These plants were used to measure photosynthesis, stomatal conductance and transpiration rate using LI-6400XT Portable Photosynthesis System (LI-COR Biosciences, Lincoln, NE, USA) These measurements were performed exactly as described in Sharma et al. (2022).

### Microarray analyses

Harvested 21-day whole plants from three independent biological replicates were used to extract total RNAs and used for microarray analyses under MIAME compliant conditions exactly as decribed in Sharma et al. (2022). The raw data and processed data were deposited in the NCBI-GEO database (GSE140257). Gene Ontology (GO) based functional annotation of differentially expressed genes (DEGs) was performed using EXPath 2.0. (http://EXPath.itps.ncku.edu.tw.) Protein subcellular localization was predicted using cropPAL database (https://crop-pal.org/) using default parameters for rice plants. MS Excel was used for filtering of the data and Student’s T-test was used for statistical significance. Venn diagrams were made using Venny 2.1 tool (https://bioinfogp.cnb.csic.es/tools/venny/).

PlantPAN3 (http://plantpan3.itps.ncku.edu.tw/) <u>was used to retrieved</u> Transcription factors (TFs) encoded by DEGs of both Nidhi and Panvel1. <u>2 kb promter regions upstream to the transcriptional initiation sites were downloaded from RAPDB and used for identification of transcription factor binding sites (TFBS) by RSAT tool</u> (http://plants.rsat.eu). Transporters encoded by DEGs were retrieved from the Rice transporters database (https://ricephylogenomics.ucdavis.edu/transporter/) and Transport DB 2.0 (http://www.membranetransport.org/transportDB2/index.html). Plant miRNA database was used to retrieve the miRNAs that target NUE-related genes (PMRD- http://bioinformatics.cau.edu.cn/PMRD/). The database Plant PTM Viewer was used to find the products of DEGs associated with post translational modifications (PTM) (https://www.psb.ugent.be/webtools/ptm-viewer/experiment.php). N-responsive yield related genes, that were defined as NUE genes, identified as described by Kumari et al. (2021) and they were co-localized to NUE-QTLs for filtering the important NUE-candidates.

### RT-qPCR validation of urea-responsive expression of DEGs

Twenty-one days old whole plants grown in normal and low urea treatments (15 mM as control and 1.5 mM as test) were used to isolate total RNAs. cDNAs were synthesized using 3 µg each of the total RNA and PrimeScript 1st strand cDNA synthesis kit (Takara, Kusatsu, Shiga, Japan). Exon spanning primes were made, using the Quant Prime tool (https://quantprime.mpimp-golm.mpg.de/?page=about) to eliminate the chances of amplifying the genomic DNA. RT-qPCR reactions were set up in an Agilent Aria-Mx Real-Time PCR exactly as described in Sharma et al. (2022). The relative changes in gene expression were quantified by 2^−ΔΔCT^ method (Livak and Schmittgen 2001) using actin genes (BGIOSGA013463) as internal controls. Melting curve analyses of the amplicons were used to determine the specificity of RT-qPCR reactions. The data were statistically analyzed by unpaired t-test using MS Excel software.

### Field experiments for agronomic, physiological and NUE

In order to test dose-specific urea responses on agronomical and physiological traits and NUE, contrasting rice genotypes Nidhi and Panvel were evaluated under field conditions at the ICAR-National Rice Research Institute, Cuttack, Odisha, India during Kharif season of 2021. Seeds were sown on 05.07.2021 and one month-old seedlings were transplanted on 10.08.2021 in a split-plot design. The plot size was 5.4 m^2^ (10 rows x 18 hills) with N levels as the main plot and varieties as the subplot in two replicates. The crop geometry was 20 cm between rows and 15 cm between plants. Urea was the sole source of applied N at the rate of 50 (N50) and 100 (N100) kg N/ha in three splits (1/2 at basal, 1/4 at vegetative, and 1/4 at flowering stage), with a control of no added N (N0). The same plots were used for N0, N50, and N100 for both the genotypes. The characteristics of the field soil before and after the experiment, along with the total carbon and nitrogen contents of plants are provided as supplementary methods.

The genotypes were harvested on 15.11.2021 at physiological maturity and various yield parameters viz., yield per panicle, grain yield per hactare, 1000 grain weight were recoreded. NUE indices were calculated as per their standard definitions as follows: Partial factor productivity (PFP, kg grain/kg N added; Grain Yield / N applied), uptake efficiency UpE = N uptake(Nt)/N supplied(Ns), nitrogen transport efficiency (NTE = total N transport into the above ground parts/total N in to the whole plants or N-uptake by plant), Utilization efficiency = grain weight/total N in plant and Fertilizer use efficiency = (fertilizer uptake/fertilizer applied)*100 were measured at maturity stage only.

## Author Contributions

NS performed most of the experiments, data analysis, and wrote the first draft. SK performed NUE-gene identification, their co-localization to NUE-QTLs, TFBS ontology and helped in primer designing, in raising and harvesting plant tissues, RNA isolation, and RT-qPCR. DKJ performed statistics of microarray, networks analyses, comparative nitrate and urea analyses, hypothetical model, and helped in final MS drafting and RT-qPCR. GKD, SP and AA did field experiments for NUE and NR helped in the planning, mentoring, and supervision of the experiments, data interpretation, and manuscript preparation. All the authors contributed to the article and approved the submitted version.

## Funding

We thank research grants from NICRA-ICAR [F. No. 2-2(60)/10-11/NICRA], DBT-NEWS-India-UK (BT/IN/UK-VNC/44/NR/2015-531 16), and UKRI-GCRF South Asian Nitrogen Hub (SANH) (NE/S009019/1), including fellowship to NS, apart from his earlier UGC-NET Fellowship.

## Conflict of Interest

The authors declare that the research was conducted in the absence of any commercial or financial relationships that could be construed as a potential conflict of interest.

## Acknowledgments

We thank Dr. Kuljeet S. Sandhu for helping in the retrieval of DEGs from raw microarray data, Vikas Kumar Mandal and Pradeep Kumar for their technical help and Dr. Padmini Swain, director of ICAR-NRRI, Cuttack for institutional support in field experiments.

## Supporting information

### Supplementary Tables

**Supplementary Table S1:** Nidhi and Panvel1 Urea DEGs

**Supplementary Table S2:** Gene Ontology analysis of Nidhi and Panvel1 DEGs

**Supplementary Table S3:** List of the primers used in this study.

**Supplementary Table S4:** Transporters of genotype Nidhi and Panvel1 encoded by DEGs, their genes, families, functions and references

**Supplementary Table S5:** Functional categorization of Transporters in Nidhi and Panvel1

**Supplementary Table S6:** Transcription Factors of genotype Nidhi and Panvel1 encoded by DEGs, their genes, families, functions and references

**Supplementary Table S7:** Functional categorization of Transcription factors in Nidhi and Panvel1

**Supplementary Table S8:** Conserved motifs in the promoter regions of Nidhi and Panvel1 urea DEGs, their TF family and gene ontology analysis

**Supplementary Table S9:** miRNAs of genotype Nidhi and Panvel1 encoded by DEGs, their target genes, functions and references

**Supplementary Table S10:** Details of protein-protein interaction (PPI) networks and their GO-based functional annotations

**Supplementary Table S11:** Details of urea-responsive PPI molecular complexes in Nidhi and Panvel1.

**Supplementary Table S12:** Details of the DEGs-associated post-translational modifications in Nidhi and Panvel1

**Supplementary Table S13:** DEGs encoding proteins for transporters and TFs

**Supplementary TableS14:** NUE-QTLs (from literature) and Nidhi urea NUE-genes colocalized onto NUE-QTLs

**Supplementary Table S15:** NUE-QTLs (from literature) and Panvel1 urea NUE-genes colocalized onto NUE-QTLs

**Supplementary Table S16:** Urea- and nitrate-responsive common and exclusive genes and associated pathways in Nidhi and Panvel1.

### Supplementary Figures

**Supplementary Figure S1:** Scatter plots show the correlation between the microarray data of Nidhi and Panvel1 grown under low (1.5 mM) and normal (15 mM) urea.

**Supplementary Figure S2:** Heatmap represents the expression pattern of urea regulated differentially expressed top two transporters and transcription factors’ families for Nidhi and Panvel1 rice contrasting genotypes.

**Supplementary Figure S3:** DEGs associated protein-protein interaction (PPI) networks developed in Nidhi under low urea condition.

**Supplementary Figure S4**: DEGs associated protein-protein interaction (PPI) networks developed in Panvel1 under low urea condition.

**Supplementary Figure S5:** DEGs associated molecular complexes/subclusters detected in the protein-protein interaction networks developed in Nidhi.

**Supplementary Figure S6:** DEGs associated molecular complexes/subclusters detected in the protein-protein interaction networks developed in Panvel1.

